# BHLHE40 drives protective polyfunctional CD4 T cell differentiation in the female reproductive tract against *Chlamydia*

**DOI:** 10.1101/2023.11.02.565369

**Authors:** Miguel A. B. Mercado, Qiang Li, Charles M. Quick, Yejin Kim, Rachel Palmer, Lu Huang, Lin-Xi Li

**Affiliations:** Department of Microbiology and Immunology, University of Arkansas for Medical Sciences, Little Rock, Arkansas 72205; Department of Pathology, University of Arkansas for Medical Sciences, Little Rock, Arkansas 72205

**Keywords:** Bhlhe40, female reproductive tract, polyfunctional CD4 T cells, Chlamydia

## Abstract

The protein basic helix-loop-helix family member e40 (BHLHE40) is a transcription factor recently emerged as a key regulator of host immunity to infections, autoimmune diseases and cancer. In this study, we investigated the role of *Bhlhe40* in protective T cell responses to the intracellular bacterium *Chlamydia* in the female reproductive tract (FRT). Mice deficient in *Bhlhe40* exhibited severe defects in their ability to control *Chlamydia muridarum* shedding from the FRT. The heightened bacterial burdens in *Bhlhe40^−/−^*mice correlated with a marked increase in IL-10-producing T regulatory type 1 (Tr1) cells and decreased polyfunctional CD4 T cells co-producing IFN-γ, IL-17A and GM-CSF. Genetic ablation of IL-10 or functional blockade of IL-10R increased CD4 T cell polyfunctionality and partially rescued the defects in bacterial control in *Bhlhe40^−/−^* mice. Using single-cell RNA sequencing coupled with TCR profiling, we detected a significant enrichment of stem-like T cell signatures in *Bhlhe40*-deficient CD4 T cells, whereas WT CD4 T cells were further down on the differentiation trajectory with distinct effector functions beyond IFN-γ production by Th1 cells. Altogether, we identified *Bhlhe40* as a key molecular driver of CD4 T cell differentiation and polyfunctional responses in the FRT against *Chlamydia*.

## Introduction

CD4 T helper cells play a central role in adaptive immunity by providing essential help to both antibody-producing B cells and cytotoxic CD8 T cells in addition to eliciting their own effector functions. Depending on the context of infection, CD4 T cells can differentiate into a variety of Th subsets (Th1/Th2/Th17/pTreg/Tfh) following antigen-specific priming by dendritic cells (Zhu et al., 2010). Mucosal infection by the obligate intracellular bacterium *Chlamydia* induces a strong type 1 immune response characterized by robust IFN-γ production by both CD4 and CD8 T cells (D. Helble and N. Starnbach, 2021). Although it is well-documented that protective immunity to *Chlamydia* relies on CD4 cells via both IFN-γ-dependent and -independent mechanisms in the female reproductive tract (FRT), the IFN-γ-independent CD4 T cell functions are largely undefined (Perry et al., 1997; Coers et al., 2011; Haldar et al., 2016; Bakshi et al., 2018; Mercado et al., 2021; Rixon et al., 2022). Moreover, the key molecules that drive the differentiation of protective CD4 T cells in *Chlamydia* infection remain to be identified.

The protein basic helix-loop-helix family, member e40 (BHLHE40) is a widely expressed transcription factor that regulates a broad range of biological processes including circadian rhythm, lipid metabolism, neurogenesis and host immunity (Kato et al., 2014; Cook et al., 2020). At steady state, *Bhlhe40* is expressed by several myeloid cell populations including neutrophils, macrophages and dendritic cells (Lin et al., 2016). In alveolar and large peritoneal macrophages, *Bhlhe40* expression is required for proliferation, self-renewal and their responses to helminth infections (Rauschmeier et al., 2019; Jarjour et al., 2019). Although not detected in naïve T cells, *Bhlhe40* expression is quickly induced upon T cell activation and maintained in a CD28-dependent manner (Martínez-Llordella et al., 2013). Recent studies describe *Bhlhe40* as a “molecular switch” between proinflammatory and anti-inflammatory responses in CD4 T cells for its dynamic regulation of cytokine production (Yu et al., 2018). Given that immune-mediated protection requires a fine balance between pro-inflammatory response that limits pathogen replication and anti-inflammatory response that prevents immunopathology, *Bhlhe40* emerges as a key player in host immune responses to infections, autoimmune disorders and cancer (Li et al., 2019; Cook et al., 2020). In *Mycobacterium tuberculosis* and *Toxoplasma gondii* infections, *Bhlhe40*^−/−^ mice manifest increased pathogen burdens and reduced survival, owing to an increase in IL-10 production by CD4 T cells and reduced IFN-γ responses in both models (Huynh et al., 2018; Yu et al., 2018). On the contrary, *Bhlhe40* deficiency ameliorates immunopathology in autoimmune conditions such as colitis and experimental autoimmune encephalomyelitis (EAE), the mouse model of multiple sclerosis (Martínez-Llordella et al., 2013; Lin et al., 2014, 2016; Yu et al., 2018). Mechanistically, BHLHE40 suppresses IL-10 production by co-binding to a regulatory region with c-Maf in the *Il10* locus in T cells and myeloid cells, which in turn promotes proinflammatory cytokine production during intracellular parasite infections. Moreover, *Bhlhe40* can also function as a transcription activator to promote *Csf2* transcription for GM-CSF production, a hallmark cytokine for pathologic CD4 T cells in EAE. Nevertheless, a role for *Bhlhe40* appears to be context dependent, as in other intracellular infection models such as *Listeria monocytogenes*, *Bhlhe40* is largely dispensable for host resistance (Huynh et al., 2018).

To date, a role for *Bhlhe40* in the FRT mucosa has not been demonstrated. Using the mouse model of *Chlamydia muridarum* intravaginal infection, we showed that loss of *Bhlhe40*, either in the germline or specifically in T cells, resulted in higher bacterial burdens and significant delay in *Chlamydia* clearance from the FRT. The defects of *Bhlhe40^−/−^* mice were results of impaired CD4 T cell responses, including increased IL-10-producing Tr1 differentiation and reduced Th1, Th17 and ThGM populations and cytokine production. Moreover, single-cell analysis revealed a fundamental defect in CD4 T cell differentiation in *Bhlhe40^−/−^* FRT manifested by increased stemness- and decreased effector function signatures in the *Bhlhe40^−/−^* CD4 T cells. These results established *Bhlhe40* as a novel regulator of CD4 T cell differentiation and protective immune responses in the FRT against *Chlamydia*.

## Results

### Bhlhe40 expression by T cells is required for anti-Chlamydia immunity

To investigate whether *Bhlhe40* plays a role in protective immunity against *Chlamydia* in the female reproductive tract (FRT), we infected WT and *Bhlhe40^−/−^* mice intravaginally with *Chlamydia muridarum* and monitored bacterial shedding from the lower FRT by vaginal swabs. As expected, WT B6 mice showed effective bacterial control and naturally resolved the infection around day 35 (**Fig. 1A-B**). In contrast, *Bhlhe40^−/−^* mice exhibited persistent high bacterial shedding from the FRT during the first 3 weeks of infection. Although bacterial control was eventually achieved in *Bhlhe40^−/−^* mice, clearance was significantly delayed (Fig. 1A-B). The differences in bacterial burden between WT and *Bhlhe40^−/−^* mice were most prominent between days 14 and 35, with 2-5 magnitudes higher inclusion forming units (IFUs) detected in *Bhlhe40^−/−^* mice than in their WT counterparts, indicating a defective adaptive immune response in these mice. On the contrary, no difference in serum antibody titers was detected on both day 21 and day 70 post infection **(Fig. S1A)**. In addition, long-term FRT tissue pathology was not affected by *Bhlhe40* deficiency, as WT and *Bhlhe40^−/−^* mice had similar pathology scores in uterine horns and oviducts at 140-150 days post infection (dpi) **(Fig. S1B)**. These data suggest that *Bhlhe40* regulates host susceptibility to *Chlamydia* infection in the FRT, likely in a T cell-dependent manner.

**Fig. 1.**
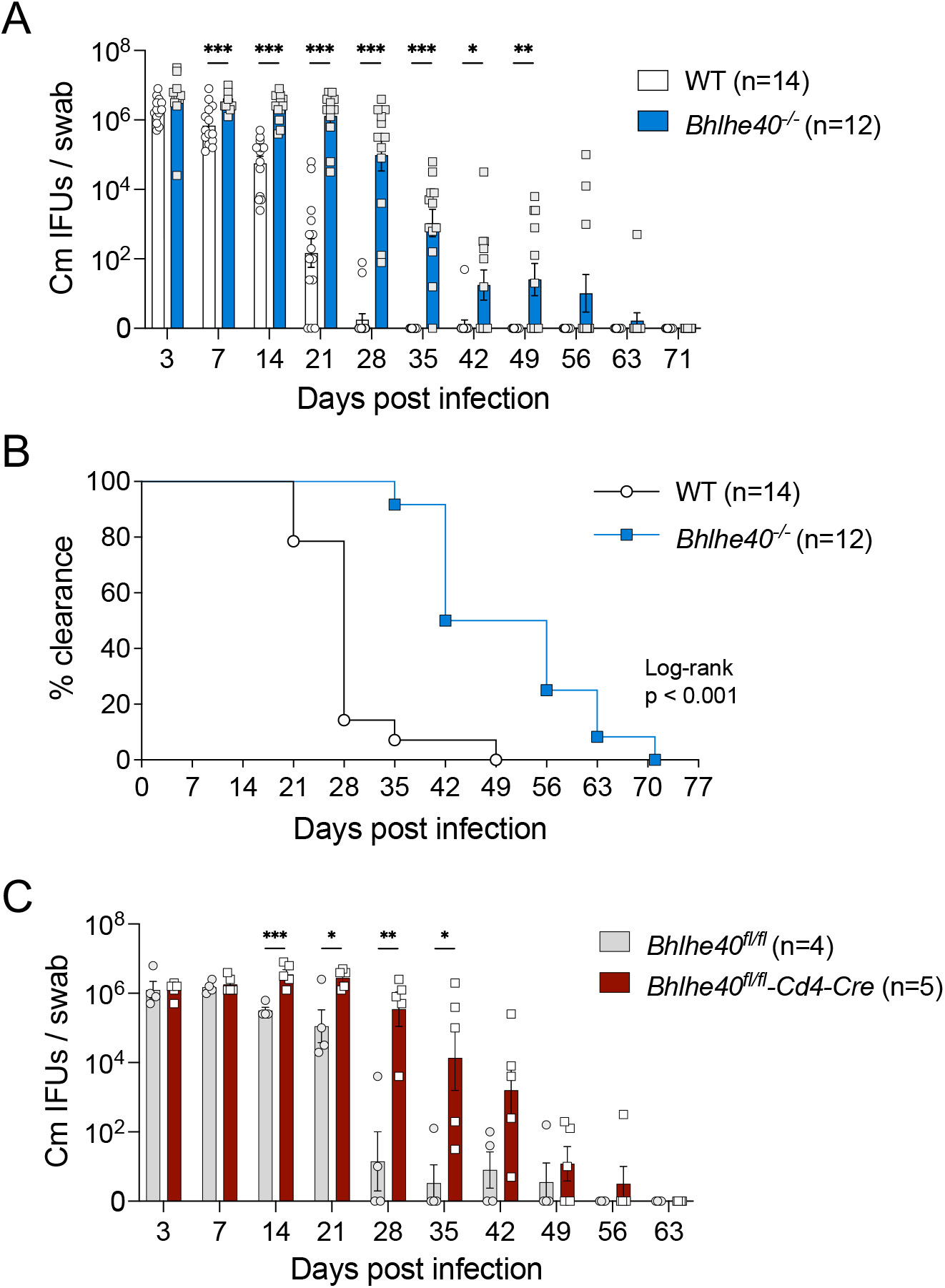
*Bhlhe40^−/−^* and *Bhlhe40^fl/fl^-Cd4-Cre* mice exhibit delayed bacterial clearance following *Chlamydia muridarum* intravaginal infection. **(A-B)** WT and *Bhlhe40^−/−^* mice were infected intravaginally with 1×10^5^ *C. muridarum*. Bacterial shedding from the lower female reproductive tract (FRT) (A) and percentage of bacterial clearance (B) were monitored by vaginal swabs. Data are combined results of three independent experiments with 12 to 14 mice per group. **(C)** *Bhlhe40^fl/fl^* and *Bhlhe40^fl/fl^-Cd4-Cre* mice were infected intravaginally with 1×10^5^ *C. muridarum*. Bacterial shedding were monitored by vaginal swabs. Data are from one experiment with 4 to 5 mice per group. Each data point represents an individual mouse. Error bars represent the mean ± SEM. *, *p <* 0.05; **, *p <* 0.01; ***, *p <* 0.001.

To determine whether T cell-intrinsic expression of *Bhlhe40* is required for anti-*Chlamydia* immunity, we next infected *Bhlhe40^fl/fl^-Cd4-Cre* mice with *C. muridarum* and compared bacterial burdens to *Bhlhe40^fl/fl^* controls. Much like the *Bhlhe40^−/−^* mice, *Bhlhe40^fl/f/^-Cd4-Cre* mice exhibited heightened bacterial shedding and delayed clearance. Moreover, the kinetics of bacterial shedding in *Bhlhe40^fl/f/^-Cd4-Cre* mice were similar to *Bhlhe40^−/−^* mice (**Fig. 1C**), indicating that T cell-intrinsic *Bhlhe40* expression accounts for the defects in bacterial control observed in *Bhlhe40^−/−^* mice.

### IL-10-producing Tr1 cells are increased in the FRT of Bhlhe40^−/−^ mice

The T cell-intrinsic requirement of *Bhlhe40* prompted us to identify the essential CD4 T cell effector functions controlled by *Bhlhe40* during *Chlamydia* infection. Previous studies have reported a disturbed balance between proinflammatory IFN-γ and anti-inflammatory IL-10 in *Bhlhe40^−/−^*mice during intracellular parasite infections (Yu et al., 2018; Huynh et al., 2018). We therefore measured CD4 T cell cytokine production in WT and *Bhlhe40^−/−^* mice at day 14 post *C. muridarum* intravaginal infection. As expected, a strong Th1 response was observed in WT mice, demonstrated by robust IFN-γ production by activated CD4 T cells (CD44^hi^) in the spleen, draining iliac lymph nodes (DLNs) and FRT (**Fig. 2A**). Although similar numbers of total and activated CD44^hi^ CD4 T cells were present in WT and *Bhlhe40^−/−^* mice (not depicted), lower frequencies and total numbers of IFN-γ-producing cells were found in the *Bhlhe40^−/−^* FRT (**Fig. 2B**). Moreover, less IFN-γ was produced on a per cell basis in IFN-γ^+^ CD4 T cells in *Bhlhe40^−/−^*mice compared to WT (**Fig. 2C**). Consistent with previous reports, we observed an increase in the frequencies of IL-10-producing cells in *Bhlhe40*-deficient mice, and IFN-γ and IL-10 double-producing CD4 T cells were also more abundant in the spleen, DLNs, and FRT of *Bhlhe40^−/−^* mice (Fig. 2B). It was evident that the increased IFN-γ^+^IL-10^+^ CD4 T cells in *Bhlhe40^−/−^* mice were T regulatory 1 (Tr1) cells rather than regulatory T cells (Tregs), as these cells do not co-express the transcription factor *Foxp3* (**Fig. S2**). These findings confirmed that *Bhlhe40* promotes the production of pro-inflammatory Th1 cells while inhibiting the differentiation of immunosuppressive Tr1 cells in the FRT.

**Fig. 2.**
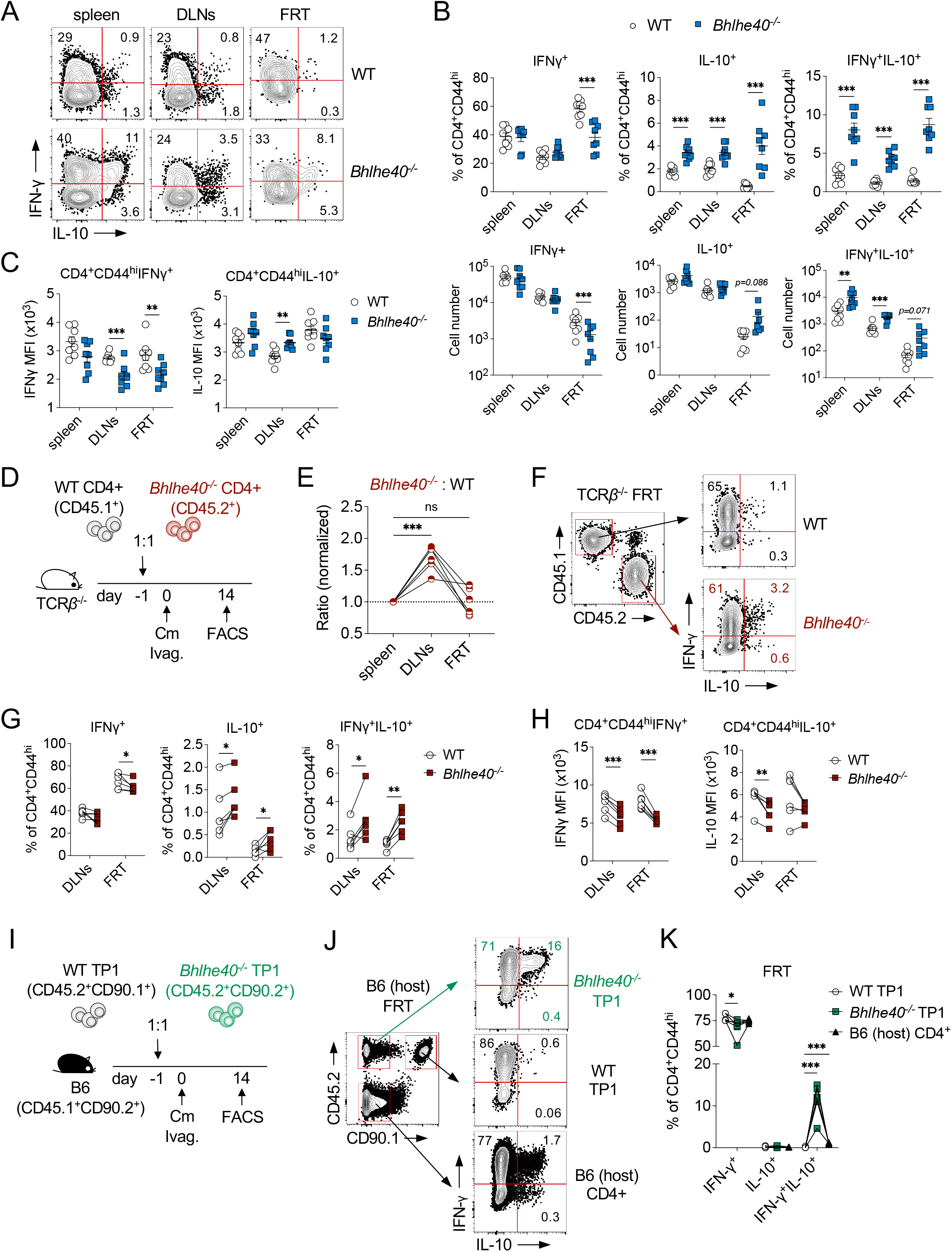
*Bhlhe40* suppresses *Chlamydia*-specific Tr1 differentiation in a T cell-intrinsic manner. **(A-C)** WT and *Bhlhe40^−/−^* mice were infected intravaginally with 1×10^5^ *C. muridarum* and analyzed at 10 dpi. (A) Representative FACS plots depicting IFNγ- and IL-10-producing CD4 T cells (gated on live CD90.2^+^CD4^+^CD44^hi^ cells). (B) Percentages and total cell numbers of IFNγ^+^, IL-10^+^ and IFNγ^+^IL-10^+^ CD4 T cells within the CD4^+^CD44^hi^ population. (C) Mean fluorescence intensities (MFIs) of IFNγ and IL-10 in CD4^+^CD44^hi^IFNγ^+^ and CD4^+^CD44^hi^IL-10^+^ T cells, respectively. Data are from two independent experiments with 8 mice per group. Each data point represents an individual mouse. **(D-H)** Mixed WT and *Bhlhe40^−/−^* CD4 T cell adoptive transfer. (D) Experimental workflow. (E) Normalized ratios of *Bhlhe40^−/−^* (CD45.2^+^) and WT (CD45.1^+^) donor CD4 T cells in TCRβ^−/−^ host. Representative FACS plots (F), summary data (G) and MFI (H) of cytokine-producing donor CD4 T cells in TCRβ^−/−^ FRT. Data are from two independent experiments with 6 mice per group. Each data point represents an individual mouse. **(I-K)** *Chlamydia*-specific TP1 CD4 T cell adoptive transfer. (I) Experimental workflow. Representative FACS plots (J) and summary data (K) of cytokine-producing *Bhlhe40^−/−^* TP1 (CD45.2^+^CD90.1^−^), WT TP1 (CD45.2^+^CD90.1^+^) and B6 host (CD45.2^−^CD90.1^−^) CD4 T cells in the host FRT. Data are from two independent experiments with 6 samples per group. Each data point represents a pooled FRT sample from 4 mice. Error bars represent the mean ± SEM. *, *p <* 0.05; **, *p <* 0.01; ***, *p <* 0.001; ns, not significant.

### Bhlhe40 suppresses antigen-specific Tr1 cell differentiation in a CD4 T cell intrinsic manner

We next sought to determine whether the skew towards anti-inflammatory Tr1 response in *Bhlhe40^−/−^* mice was due to cell-intrinsic regulation of CD4 T cell differentiation by *Bhlhe40*. To do this, we co-transferred WT (CD45.1^+^) and *Bhlhe40*-deficient (CD45.2^+^) naïve CD4 T cells into TCRβ^−/−^ mice at a 1:1 ratio and infected the recipients intravaginally with *C. muridarum* (**Fig. 2D**). At 14 dpi, higher percentages of *Bhlhe40^−/−^*CD4 T cells were observed in the host DLNs (normalized ratio of *Bhlhe40^−/−^*: WT = 1.62 ± 0.26) (**Fig. 2E**), presumably due to enhanced T follicular helper cell proliferation within the *Bhlhe40^−/−^* CD4 T cell population (Rauschmeier et al., 2021). In contrast, comparable numbers of WT and *Bhlhe40^−/−^* CD4 T cells were found in the FRT, indicating that *Bhlhe40^−/−^* CD4 T cells had no defect in tissue homing or accumulation in this mucosal site. When quantifying cytokine producing cells within each donor compartment, we found that the frequencies of IFN-γ^+^ cells were slightly lower in the *Bhlhe40^−/−^* T cell compartment in the FRT, and IFN-γ production on a per cell basis was significantly lower in *Bhlhe40*-deficient CD4 T cells in both DLNs and FRT compared to WT (**Fig. 2F-H**). Consistent with our findings in *Bhlhe40^−/−^* mice, the frequencies of IL-10^+^ CD4 T cells and IFN-γ^+^IL-10^+^ Tr1 cells were significantly higher in the *Bhlhe40^−/−^* CD4 T cell compartment in the DLNs and FRT (Fig. 2G). These results demonstrated that although *Bhlhe40* is widely expressed in many leukocyte lineages, it suppresses Tr1 differentiation in a CD4 T cell-intrinsic manner independent of other cell types or environment cytokine milieu.

*Chlamydia* infection in the FRT drives a potent CD4 T cell expansion in the secondary lymphoid organs and recruits both antigen-specific and bystander CD4 T cells to the site of infection (Li and McSorley, 2013; O’Donnell et al., 2014). These two populations contribute differently to host protective immunity and development of immunopathology (McSorley, 2014; Lijek et al., 2018). To interrogate the role of *Bhlhe40* in antigen-specific CD4 T cells, we crossed the *Chlamydia*-specific TCR transgenic mice (TP1) onto the *Bhlhe40*-deficient background and conducted co-transfer experiments with WT (CD45.2^+^CD90.1^+^) and *Bhlhe40*-deficient (CD45.2^+^CD90.2^+^) TP1 cells (**Fig. 2I**) (Poston et al., 2017). Using CD45.1^+^ B6 mice as recipients, we were able to detect WT TP1, *Bhlhe40^−/−^* TP1 and endogenous CD4 T cells within the same host (**Fig. 2J**). At 14 dpi, WT TP1 cells were predominantly IFN-γ^+^ cells with minimum IL-10^+^ CD4 T cells or IFN-γ^+^IL-10^+^ Tr1 cells detected (**Fig. 2J and 2K**). Endogenous CD4 T cells contained comparable frequencies of IFN-γ^+^ cells to WT TP1, with a small but evident Tr1 cell population also detected in this population. Notably, *Bhlhe40^−/−^*TP1 cells contained the highest percentages of Tr1 cells with lower frequencies of IFN-γ single-producers compared to WT TP1 (Fig. 2J and 2K). Thus, we conclude that *Bhlhe40* is essential for driving Tr1 differentiation in antigen-specific CD4 T cells during *Chlamydia* FRT infection.

### Increased IL-10-producing Tr1 cells dampen protective CD4 T cell responses in Bhlhe40^−/−^ mice

We next aimed to test whether elevated IL-10 production by increased immunosuppressive Tr1 cells in *Bhlhe40^−/−^* mice accounts for the defects in bacterial control in these mice. WT and *Bhlhe40^−/−^* mice were treated with either Anti-IL-10R or isotype control antibody starting 1 day prior to *C. muridarum* intravaginal infection. As shown in **Fig. 3**, IL-10R blockade significantly reduced bacterial shedding in both WT and *Bhlhe40^−/−^* mice compared to isotype-treated groups, confirming the immunosuppressive function of IL-10 in both mouse models. Moreover, anti-IL-10R treated WT mice mirrored the phenotype of *Il10^−/−^* mice, indicating that antibody-mediated IL-10R blockade has similar biological effects as IL-10 cytokine knockout in this infection model. Importantly, whereas anti-IL-10R treatment in *Bhlhe40^−/−^* mice reduced bacterial burdens to comparable levels as WT isotype-treated group, it did not attain the accelerated bacterial control as in WT anti-IL-10R group or *Il10^−/−^* mice. These results indicated that in addition to restricting Tr1 differentiation and IL-10 production, *Bhlhe40* modulates other CD4 T cell effector functions to facilitate *Chlamydia* clearance from the FRT.

**Fig. 3.**
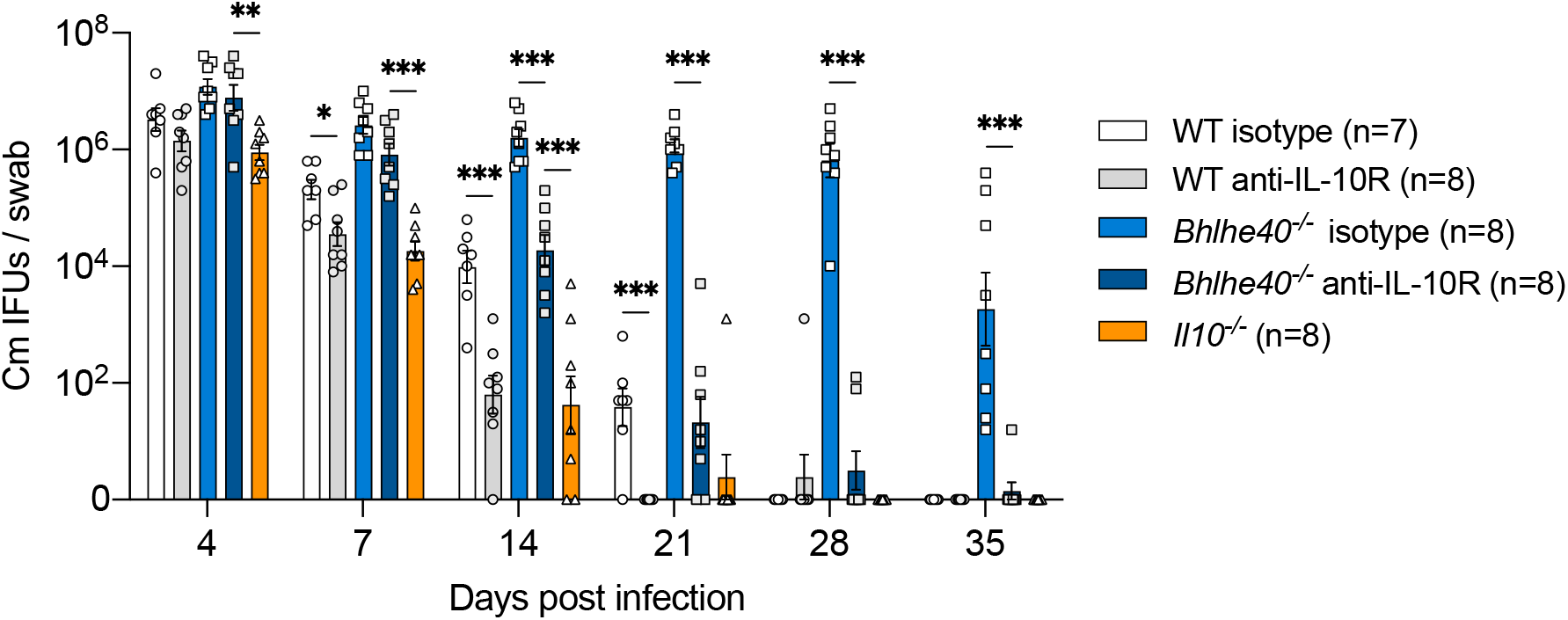
Anti-IL-10R blockade partially restores the ability of bacterial control in *Bhlhe40*-deficient mice. WT, *Bhlhe40^−/−^* and *Il10*^−/−^ mice were infected intravaginally with 1×10^5^ *C. muridarum*. Cohorts of WT and *Bhlhe40^−/−^* mice were treated with anti-IL-10R or isotype control antibody twice a week until day 21. Bacterial shedding from the lower FRT were monitored by vaginal swabs. Data are results from two independent experiments with 7-8 mice per group. Each data point represents an individual mouse. Error bars represent the mean ± SEM. *, *p <* 0.05; **, *p <* 0.01; ***, *p <* 0.001.

### Bhlhe40 promotes the differentiation of protective polyfunctional CD4 T cells

To expand our investigation of how *Bhlhe40* regulates CD4 T cell effector functions, we performed a multiplex cytokine array of ex vivo cultures of DLNs and FRT cells isolated from *C. muridarum* infected WT and *Bhlhe40^−/−^* mice. At 10 dpi, major Th1 cytokines, including IFN-γ, TNF-α and IL-2, were significantly lower in the *Bhlhe40^−/−^* FRT compared to WT (**Fig. 4A, Table S1**). Moreover, IL-17A and GM-CSF, signature cytokines produced by Th17 and ThGM cells, respectively, were also drastically reduced in the absence of *Bhlhe40*. These observations were reinforced by intracellular cytokine staining in which lower percentages of IL-17A^+^ and GM-CSF^+^ CD4 T cells were detected in *Bhlhe40^−/−^* mice (**Fig. 4B**), suggesting that *Bhlhe40* regulates a broader range of CD4 T cell effector functions beyond Th1 and Tr1 differentiation. It is worth noting that although IL-10-producing Tr1 cells were increased in *Bhlhe40^−/−^* mice, we did not detect a difference in IL-10 levels in this ex vivo cell culture setting (Fig. 4A). We next sought to quantify CD4 T cells capable of producing more than one pro-inflammatory cytokine. SPICE analysis showed that WT mice contained higher proportions of CD4 T cells capable of producing 2 or 3 cytokines than *Bhlhe40^−/−^*mice (**Fig. 4C**). Within WT polyfunctional CD4 T cells, IFN-γ^+^GM-CSF^+^ cells were the most abundant, followed by IFN-γ^+^IL-17A^+^ and IFN-γ^+^IL-17A^+^GM-CSF^+^ cells (**Fig. 4D**). To ask whether this phenotype recapitulates in *Chlamydia*-specific CD4 T cells, we utilized the TP1 mixed transfer strategy described above (Fig. 2I) and measured cytokine productions from both TCR Tg and endogenous CD4 T cells. Interestingly, unlike polyclonal CD4 T cells in *Bhlhe40^−/−^* mice, *Bhlhe40^−/−^* TP1 cells contained higher frequencies of CD4 T cells capable of producing IL-17A, as percentages of both IL-17A^+^ and IFN-γ^+^IL-17A^+^ cells were considerably higher in *Bhlhe40^−/−^* TP1 compared to WT TP1 or endogenous CD4 T cells (**Fig. 4E and 4F**). In contrast, lower frequencies of IFN-γ^+^GM-CSF^+^ cells were detected in *Bhlhe40^−/−^* TP1 cells. The overall CD4 T cell polyfunctionality was comparable between WT and *Bhlhe40^−/−^* TP1 cells (not depicted). Together, these data suggest that *Bhlhe40* positively regulates T cell GM-CSF production, and potentiates polyfunctional CD4 T cell differentiation via a combined T cell-intrinsic and -extrinsic mechanism.

**Fig. 4.**
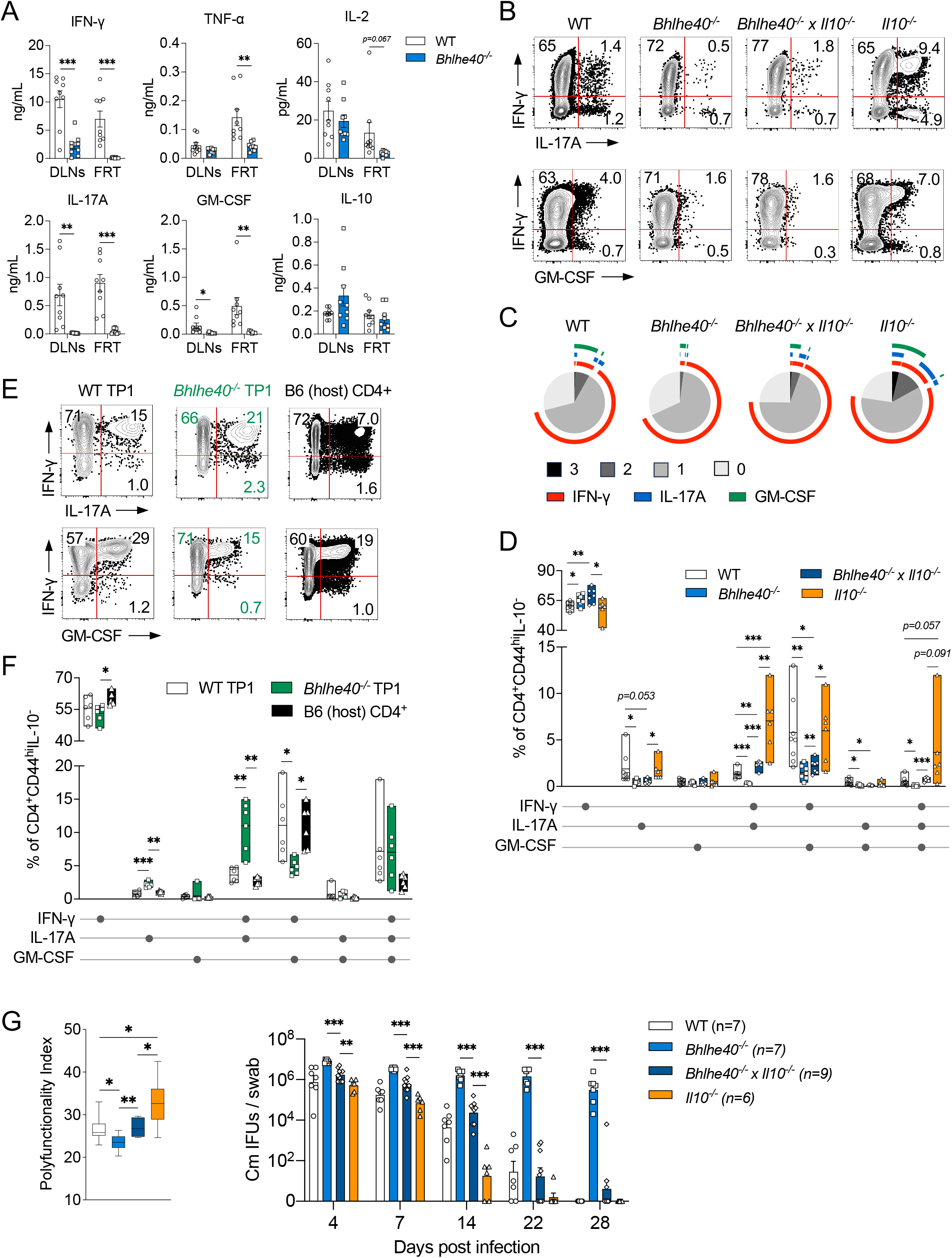
CD4 T cell polyfunctionality inversely correlates with host susceptibility to *C. muridarum* in the FRT. **(A)** WT and *Bhlhe40^−/−^* mice were infected intravaginally with 1×10^5^ *C. muridarum*. Cytokine levels in the supernatants of *ex vivo* cultures of DLN and FRT cells as measured at 10 dpi. Data are from two independent experiments with 9 mice per group. Each data point represents an individual mouse. **(B-D)** WT, *Bhlhe40^−/−^*, *Bhlhe40^−/−^* x *Il10*^−/−^ and *Il10*^−/−^ mice were infected intravaginally with 1×10^5^ *C. muridarum*. Representative FACS plots (B), SPICE analysis (C) and summary data (D) depicting cytokine producing CD4 T cells (gated on live CD90.2^+^CD4^+^CD44^hi^IL-10^−^ cells) at 14 dpi. Data are from two independent experiments with 6-9 mice per group. Each data point represents an individual mouse. **(E-F)** *Chlamydia*-specific TP1 CD4 T cell adoptive transfer experiment (see workflow in Fig. 2I). Representative FACS plots (E) and summary data (F) of cytokine-producing *Bhlhe40^−/−^* TP1 (CD45.2^+^CD90.1^−^), WT TP1 (CD45.2^+^CD90.1^+^) and B6 host CD4 T cells (CD45.2^−^CD90.1^−^) in the host FRT. Data are from two independent experiments with 6 samples per group. Each data point represents a pooled FRT sample from 4 mice. **(G)** Polyfunctionality Index (left) and bacterial burdens (right) in WT, *Bhlhe40^−/−^*, *Bhlhe40^−/−^* x *Il10*^−/−^ and *Il10*^−/−^ mice. Data are from two independent experiments with 6-9 mice per group. Each data point represents an individual mouse. Error bars represent the mean ± SEM. *, *p <* 0.05; **, *p <* 0.01; ***, *p <* 0.001.

It is possible that the discrepant T cell cytokine profiles in *Bhlhe40*-deficient mice and *Bhlhe40^−/−^* TP1 cells were due to the immunosuppressive environment created by increased Tr1 cell and IL-10 production in *Bhlhe40^−/−^*mice. To answer whether the Tr1/IL-10 axis curtails T cell polyfunctionality, we crossed *Bhlhe40^−/−^*mice to *Il10^−/−^* mice and infected these double knockout mice with *C. muridarum*. As expected, *Bhlhe40^−/−^* x *Il10^−/−^* and *Il10^−/−^* mice had higher percentages of polyfunctional CD4 T cells compare to *Bhlhe40^−/−^* and WT mice, respectively (Fig. 4B and 4C). Importantly, although IL-10 deficiency led to an increase in IL-17A-producing polyfunctional CD4 T cells in both WT and *Bhlhe40^−/−^* mice, it did not impose a detectable effect on IFN-γ, IL-17A, or GM-CSF single-producing T cells, and had a marginal effect on polyfunctional CD4 T cells that produce GM-CSF. These results suggest that the Tr1/IL-10 axis partially accounts for the limited polyfunctional CD4 T cell differentiation in *Bhlhe40^−/−^*mice.

Polyfunctional CD4 and CD8 T cells are associated with enhanced protective immunity during infections and vaccinations (Seder et al., 2008; Appay et al., 2008; Boyd et al., 2015). We therefore interrogated the relationship between CD4 T cell polyfunctionality and protective immunity against *Chlamydia*. As shown in **Fig. 4G**, CD4 T cell polyfunctionality index was lowest in *Bhlhe40^−/−^* mice and highest in *Il10^−/−^* mice, and these indices reversely correlated with bacterial shedding over the course of *C. muridarum* primary infection (Larsen et al., 2012). Thus, we conclude that CD4 T cell polyfunctionality is positively associated with protective immunity against *Chlamydia*.

### Single-cell RNA sequencing reveal an increase in CD4 T cell stemness in Bhlhe40^−/−^ mice

To further expand our understanding of how *Bhlhe40* regulates anti-*Chlamydia* immunity, we performed 5’ single-cell RNA sequencing (scRNAseq) and TCR profiling on activated CD4 T cells (CD44^hi^) sorted from WT and *Bhlhe40^−/−^* mouse FRT 14 days post intravaginal infection. Unsupervised clustering revealed 14 clusters from a combined population of 5,782 WT and 3,732 *Bhlhe40^−/−^* CD4 T cells (**Fig. 5A**). Following cell number normalization, we found that WT CD4 T cells were more enriched in clusters 0, 4, and 8 whereas *Bhlhe40^−/−^* CD4 T cells were dominant in clusters 3, 6 and 10 (>80% of total, **Fig. 5B**). As expected, CD4 T cells in the largest WT cluster 0 expressed high levels of *Bhlhe40*, along with a panel of genes downstream of TCR signaling including *Ifitm1*, *Ifitm2*, and *Nr4a1* (**Fig. 5C**) (Yánez et al., 2020; Osborne et al., 1994). In contrast, the predominant *Bhlhe40^−/−^*CD4 T cells clusters 3 expressed genes that resembled T cell stemness, such as *Tcf7*, *Ccr7*, *Slamf6* and *Sell* (Utzschneider et al., 2016; Nish et al., 2017). Gene expression pattern in cluster 10 closely resembled that of cluster 3, with significantly more ribosomal genes upregulated in this cluster (e.g. *Rps 29*, *Rpl12*), indicating increased protein translation. Analysis of T helper lineage specific genes revealed that majority of the clusters expressed the Th1-specific *Tbx21* and *Ifng* genes, with the lowest expression detected in *Bhlhe40^−/−^*-dominant clusters 3 and 10 (**Fig. 5D**). On top of the strong type 1 response, Th17-specific transcripts *Rorc* and *Il17a* were detected at low levels in the WT-dominant cluster 4, and this cluster also uniquely expressed Th17-related genes, including *Tmem176b*, *Ltb4r1*, *Il17re*, and *Ramp1* (Ciofani et al., 2012; Chang et al., 2011; Mikami et al., 2012), revealing its identity as polyfunctional Th1/Th17 cells. Additionally, WT cluster 8 expressed a panel of genes associated with cytotoxicity and terminal differentiation, such as *Gzma*, *Klrg1*, *Cx3cr1* and *Zeb2* (Krueger et al., 2021), indicating their cellular identify of cytotoxic CD4 T helpers. Lastly, *Bhlhe40^−/−^*cluster 6 presented a terminal-differentiated Tr1 phenotype by expressing *Lag3*, *Maf*, *Il10ra* and *Havcr2*, supporting our previous observations that Tr1 cells were increased in these mice (Roncarolo et al., 2018; Brockmann et al., 2017).

**Fig. 5.**
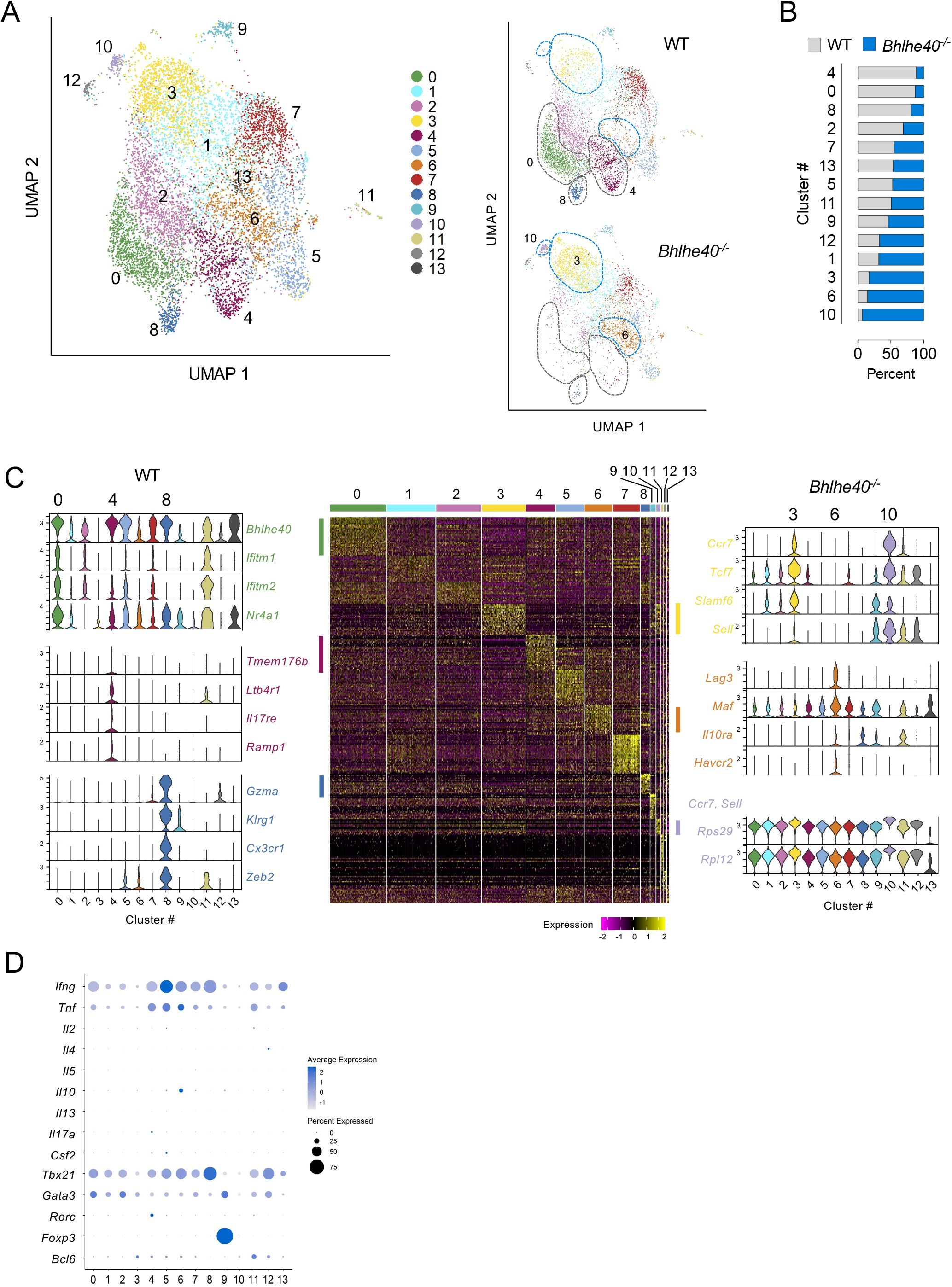
Single-cell RNA sequencing detects stem-like CD4 T cells in *Bhlhe40^−/−^* FRT. WT and *Bhlhe40^−/−^* mice were infected intravaginally with 1×10^5^ *C. muridarum*. Activated CD4 T cells (live CD90.2^+^CD4^+^CD44^hi^) in the FRT were sorted for scRNAseq using 10x genomics 5’ GEX. **(A)** UMAP depicting 14 clusters of CD4 T cells from merged WT and *Bhlhe40^−/−^* CD4 T cells (left); deconvolution of the composite UMAP into WT and *Bhlhe40^−/−^* components. **(B)** Percentages of WT and *Bhlhe40^−/−^* CD4 T cells in each cluster following normalization of the two populations into the same cell numbers. **(C)** Heatmap showing top 30 most upregulated genes in each cluster, and representative genes specifically expressed within each of the top 3 clusters in WT and *Bhlhe40^−/−^* CD4 T cells are shown on the sides with violin plots depicting expression levels in each cluster. **(D)** Dot plots depicting Th lineage-specific cytokines and transcription factors.

To delineate the cellular differentiation trajectory within these T cell clusters, we conducted pseudotime trajectory analysis using the stem-like cluster 3 T cells as origin. Most of *Bhlhe40^−/−^* T cells scored low on the differentiation trajectory, including the Tr1 cluster 6 despite the expression of several exhaustion markers by these T cells (**Fig. 6A**). In contrast, WT enriched clusters 0, 4 and 8 were among the clusters with highest differentiation scores. Clusters with intermediate differentiation scores, including clusters 2 (migratory T) and 7 (ISG-enriched T), were also over-represented by WT CD4 T cells (Fig. 6A and 5B, Table S2). To determine whether T cell clonal expansion correlates with differentiation, we analyzed TCR clonotypes in WT and *Bhlhe40^−/−^* T cells and projected the top 25% abundant clones (“expanders”) onto the T cell clusters. Similar profiles of clonal expansion were detected in WT and *Bhlhe40^−/−^* CD4 T cells, with the top 25% cell population falling between clonal abundance of 3 to 164 (**Fig. 6B**). The most differentiated clusters 0 and 8 contained highly expanded CD4 T cells, indicating that these T cells were the most dynamic populations during the immune response. Curiously, although cluster 6 (Tr1) had a less differentiated profile on the trajectory, it contained high frequencies of “expanders” in both WT and *Bhlhe40^−/−^* T cells. Altogether, these data demonstrated that *Bhlhe40* is essential for driving CD4 T cell differentiation from stem-like CD4 T cell progenitors into multi-functional effectors.

**Fig. 6.**
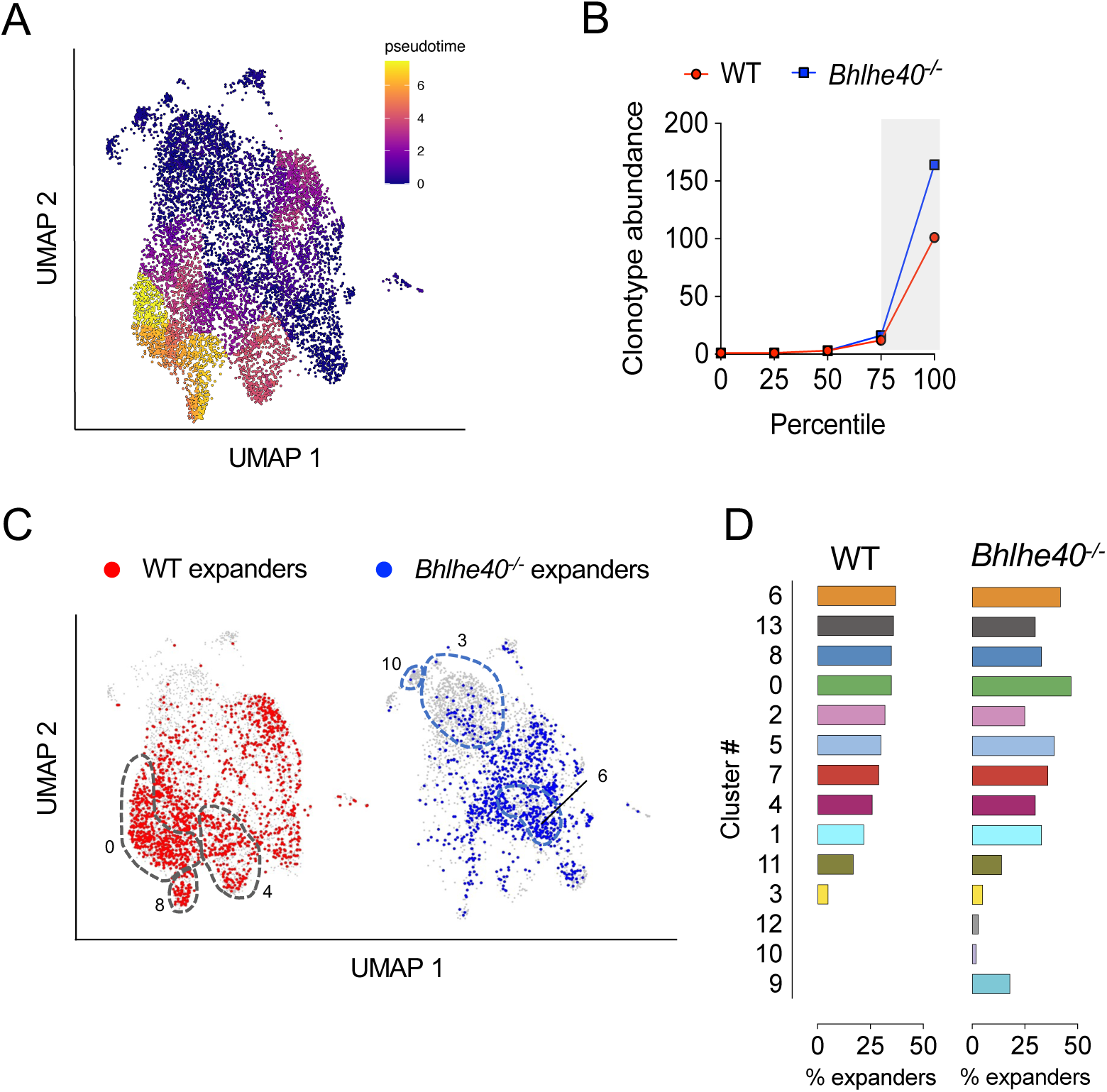
Pseudotime trajectory and TCR clonal type analyses reveal CD4 T cell differentiation and clonal expansion profiles in WT and *Bhlhe40^−/−^* mice. (A) Pseudotime trajectory analysis of CD4 T cell clusters in Fig. 5 as conducted using Monocle 3. **(B)** WT and *Bhlhe40^−/−^* CD4 T cells were binned into quartiles based on clonotype abundance. Shaded area represents the top 25% (expanders). **(C)** UMAP showing the expanders in WT and *Bhlhe40^−/−^* CD4 T cell clusters. **(D)** Frequencies of expanders in each cluster in WT and *Bhlhe40^−/−^* CD4 T cells.

## Discussion

The prevailing paradigm in protective adaptive immunity to the intracellular bacterium *Chlamydia* is centered on IFN-γ production by CD4 Th1 cells. Nevertheless, mice lacking either Th1-specific transcription factor T-bet or IFN-γ-producing CD4 T cells were both capable of restricting *Chlamydia* replication in the female reproductive tract (Mercado et al., 2021; Rixon et al., 2022). Moreover, increasing evidence suggest that multiple Th lineages and T cell effector functions, including Th2, Th17, IL-13 production and CD4 T cell-dependent cytotoxicity, may participate in host defense against *Chlamydia* (Scurlock et al., 2011; Miguel et al., 2013; Li and McSorley, 2013; Johnson et al., 2018; Labuda and McSorley, 2018). Additionally, our evolving understanding of CD4 T cell heterogeneity over recent years indicate that the functional state of CD4 T cells is likely a continuum shaped by the microbes and the tissue microenvironment rather than a defined Th subtype (Cano-Gamez et al., 2020; Kiner et al., 2021). In this current study, we provide new evidence supporting this scenario by showing that protective immunity to *Chlamydia* relies on highly differentiated polyfunctional CD4 T cells driven by the transcription factor *Bhlhe40* rather than a single Th subset or cytokine in the FRT mucosa.

Although *Bhlhe40* is widely expressed in many cell types and tissues, our data demonstrated that T cell-intrinsic expression of *Bhlhe40* is essential for *Chlamydia* control in the FRT, as *Bhlhe40*^−/−^ and *Bhlhe40^fl/fl^-Cd4-Cre* mice exhibit almost identical kinetics in bacterial clearance. Previous studies have shown that *Bhlhe40* deficiency resulted in reduced IL-2 production by T cells, and consequently decreased T cell proliferation and survival in an autocrine manner (Martínez-Llordella et al., 2013). Nevertheless, we did not observe a reduction in either total or activated CD4 T cell numbers in the *Bhlhe40*^−/−^ FRT, albeit a decreased IL-2 production was observed in *in vitro* culture assay. CD4 T cell accumulation was also unaffected by *Bhlhe40* deficiency in a competitive setting where WT and *Bhlhe40*^−/−^ CD4 T cells were co-transferred into T cell deficient TCRβ^−/−^ mice. These results collectively suggest that T cell effector functions, rather than T cell homing or accumulation at the FRT mucosa, accounts for the defects in bacterial control in the *Bhlhe40*-deficient mice during *Chlamydia* infection.

Previous studies have established a role for *Bhlhe40* in host resistance to several intracellular pathogens including *M. tuberculosis, T. gondii,* and influenza (Huynh et al., 2018; Yu et al., 2018; Li et al., 2019). Our results add to that by showing the importance for *Bhlhe40* for anti-*Chlamydia* immunity at the FRT mucosa. The defects in pathogen control in *Bhlhe40*^−/−^ mice were caused, at least in part, by increased anti-inflammatory IL-10 production from CD4^+^ Tr1 cells, as the phenotype can be partially reversed by either genetic ablation of IL-10 or antibody-mediated IL-10R blockade. Curiously, although IL-10-producing cells were increased in *Bhlhe40*^−/−^ mice, we did not observe any difference in pathological outcomes between WT and *Bhlhe40^−/−^*FRT, presumably because the increase in anti-inflammatory responses was counterbalanced by longer duration of bacterial shedding in these mice.

In both lung and FRT *Chlamydia* infection models, IL-10-deficiency leads to accelerated bacterial clearance (Yang et al., 1999; Igietseme et al., 2000). Our study recapitulates these findings and further demonstrates that IL-10 functions as a roadblock in polyfunctional CD4 T cell differentiation, as IL-10 ablation in both WT and *Bhlhe40^−/−^* mice increases T cell polyfunctionality index. The importance of multifunctional Th1 cells in protective immunity to *Chlamydia* has been demonstrated previously. Intranasal immunization with live *Chlamydia* elementary body (EB) renders better protection against *Chlamydia* FRT challenge than immunization with dead EB, and the IFN-γ^+^ TNF-α^+^IL-2^+^ triple- and IFN-γ^+^ TNF-α^+^ double-positive CD4 T cell profiles correlate with protection (Yu et al., 2011). Using TCR transgenic mice, Poston et al showed that *Chlamydia*-specific CD4 T cells preferentially adopt a polyfunctional Th1 phenotype and confer better protection at low cell numbers compared to polyclonal T cells (Poston et al., 2017). Here we expand the scope beyond the Th1 spectrum and illustrate that the frequency of polyfunctional CD4 T cells producing multiple proinflammatory cytokines, including IFN-γ, IL-17A and GM-CSF, correlates with better protective immunity to *Chlamydia*. We propose two potential interpretations of these observations: First, synergistic effects of these cytokines may be essential for protection. It has been known for some time that mice deficient in a single cytokine, such as IFN-γ or IL-17A had only minor defects in *Chlamydia* control in the FRT mucosa (Perry et al., 1997; Cotter et al., 1997; Scurlock et al., 2011; Andrew et al., 2013). The observations that *Bhlhe40*^−/−^ mice had comparable numbers of IFN-γ single-producing CD4 T cells to WT, but failed to control *Chlamydia* effectively further reinforce the notion that IFN-γ alone does not account in full for anti-*Chlamydia* immunity in the FRT. *Csf2^−/−^*mice are susceptible to several intracellular bacterial and viral infections in the lung (LeVine et al., 1999; Gonzalez-Juarrero et al., 2005; Schneider et al., 2014), but a role for GM-CSF in CD4 T cell-mediated immunity to *Chlamydia* is yet to be established. Previous studies have identified *Chlamydia*-specific IFN-γ^+^IL-17A^+^ CD4 T cells in the FRT following *Chlamydia* infection or vaccination (Li and McSorley, 2013; Nguyen et al., 2020). It is somewhat surprising that we detected IFN-γ^+^GM-CSF^+^ cells as the most abundant polyfunctional CD4 T cells in the FRT. Given that IFN-γ-producing Th1 cells can gain GM-CSF signature, IL-17A^+^ CD4 T cells can differentiate into IFN-γ^+^GM-CSF^+^ pathologic Th17 cells upon IL-23 stimulation, and a separate ThGM subset has also been proposed, the origins of IFN-γ^+^GM-CSF^+^ double- and IFN-γ^+^IL-17A^+^GM-CSF^+^ triple-producing cells in *Chlamydia* infection remain unknown (Grifka-Walk et al., 2015; Schnell et al., 2021; Zhang et al., 2013). Future studies will be needed to identify the cell fate and determine the effector functions of these populations. It is worth noting that in our model both IFN-γ^+^IL-10^+^ Tr1 and GM-CSF-producing CD4 T cells are regulated by *Bhlhe40* in a CD4 T cell-intrinsic manner, likely at the transcription level as reported before. In contrast, the reduction of IL-17A^+^ and IFN-γ^+^IL-17A^+^ CD4 T cells in *Bhlhe40*^−/−^ mice is attributed to a T cell-extrinsic factor, as the defects were not observed when *Bhlhe40*-deficient TP1 cells were primed in the WT B6 host (Fig. 4E). It would be of interest to identify the contributing cell types and/or the environmental milieu that block the differentiation of IL-17A-producing CD4 T cells in *Bhlhe40*^−/−^ mice. A second and perhaps more appealing interpretation would be that the polyfunctionality of CD4 T cells co-producing 2 or 3 cytokines simply reflects a state of differentiation required for protective immunity. This argument is supported by our scRNAseq data that *Bhlhe40*^−/−^ CD4 T cells are enriched in clusters expressing more “stem-like” T cell markers, whereas WT CD4 T cells are further down on the differentiation trajectory with diverse effector functions, along with a more robust clonal expansion profile. In line with our findings, recent work from Wherry and colleagues reported that during chronic LCMV infection, *Bhlhe40* drives exhausted CD8 T cells (T_ex_) from stem-like T_ex_ into terminal differentiated T_ex_, demonstrating its role in regulating CD8 T cell differentiation (Wu et al., 2023). Although it is difficult to precisely project each polyfunctional CD4 T cell population identified by flow cytometry to specific T cell clusters in scRNAseq due to the low transcript numbers of *Il17a* and *Csf2*, and potential discrepancy between cytokine transcription and translation, the fact that all WT clusters contain *Ifng* transcripts while co-express Th17 or cytotoxic T cell signature genes reveals polyfunctionality of these cells. The cytotoxic CD4 T cell cluster is of particular interest, as previous studies by Johnson and colleagues have showed that CD4 T cell clones had direct killing capability of infected epithelium cells *in vitro* (Jayarapu et al., 2010). The emerging roles of cytotoxic CD4 T cells have been demonstrated in a variety of infection models, autoimmune diseases and cancer (Cenerenti et al., 2022; Oh and Fong, 2021). Experiments are currently underway to evaluate whether *Chlamydia* infection induces cytotoxic function in the polyfunctional CD4 T cells *in vivo* and whether these cells contribute to protective immunity.

In summary, our study provides compelling evidence that a multifaceted CD4 T cell response in *Chlamydia* infection is essential for protective immunity. Moreover, we identify the transcription factor *Bhlhe40* as a novel regulator of CD4 T cell differentiation in the FRT. Understanding the protective features and identifying the key regulators of CD4 T cell responses provide important guidelines for future *Chlamydia* vaccine design.

## Materials and Methods

### Mice

C57BL/6 (B6), CD45.1^+^ (B6.SJL-*Ptprc^a^ Pepc^b^*/BoyJ), *Bhlhe40*^−/−^ (B6.129S1(Cg)-*Bhlhe40^tm1.1Rhli^*/MpmJ), *Il10^−/−^* (B6.129P2-*Il10^tm1Cgn^*/J) and TCRβ^−/−^ (B6.129P2-*Tcrb^tm1Mom^*/J) mice were purchased from The Jackson Laboratory (Bar Harbor, ME). *Bhlhe40^fl/fl^* mice were provided by Dr. Brian Edelson (Wash U) via Dr. Jason Stumhofer (UAMS) (Huynh et al., 2018; O’Neal et al., 2023). *Bhlhe40^fl/fl^* mice were crossed with *Cd4-cre* (B6.Cg-Tg(Cd4-cre)1Cwi/BfluJ) mice to generate *Bhlhe40^fl/fl^-Cd4-cre* mice (by Dr. Jason Stumhofer, UAMS). *Bhlhe40*^−/−^ x IL-10^−/−^ mice were generated in-house by crossing *Bhlhe40*^−/−^ mice with *Il10^−/−^* mice. *Chlamydia-*specific TCR-transgenic (TP1) mice were provided by Drs. Taylor Poston and Toni Darville (UNC Chapel Hill) (Poston et al., 2017). TP1 mice were crossed with *Bhlhe40^−/−^* mice to generate TP1 x *Bhlhe40^−/−^* mice. All mice used for experiments were 6 to 24 weeks old. Mice were maintained under SPF conditions and all mouse experiments were approved by the University of Arkansas for Medical Sciences Institutional Animal Care and Use Committee (IACUC).

### Chlamydia strain, mouse infection and bacteria enumeration

*Chlamydia muridarum* strain Nigg II was originally purchased from ATCC (VR-123; Manassas, VA). *C. muridarum* was propagated in McCoy cells, elementary bodies (EBs) purified by discontinuous density gradient centrifugation and titrated on HeLa 229 cells as previously described (Li and McSorley, 2013). Mice were synchronized for estrous by subcutaneous injection of 2.5 mg medroxyprogesterone (Depo-Provera, Greenstone, NJ) 5-7 days prior to intravaginal infection. For intravaginal (i.vag.) infection, 1×10^5^ *C. muridarum* in SPG buffer was deposited directly into the vaginal vault using a pipet tip. To enumerate bacterial shedding from the lower FRT, vaginal swabs were collected, suspended in SPG buffer, and disrupted with glass beads. Inclusion forming units (IFUs) were determined by plating serial dilutions of swab samples on HeLa 229 cells, staining with anti-MOMP mAb (clone Mo33b), and counting under a microscope.

### Histopathology

Female reproductive tracts from WT and *Bhlhe40^−/−^* mice were harvested between 140-150 days post infection and fixed in 4% paraformaldehyde overnight. The tissues were then embedded in paraffin blocks, longitudinal sections prepared and stained with hematoxylin and eosin. The sections were scored using a previously elaborated semiquantitative scoring system by a board-certified pathologist who was blind to the experimental design (Darville et al., 1997; Li et al., 2017). The following parameters were evaluated on a scale of 0 (normal) to 4 (severe lesion): acute inflammation, chronic inflammation, plasma cells, epithelial erosion, and fibrosis. Left and right sides of the uterine horns and oviducts were evaluated individually.

### Chlamydia-specific serum Ab ELISA

Mice were bled via the submandibular vein at 21 days post infection to isolate serum. Heat-killed *C. muridarum* elementary bodies (HKEBs) were prepared by heating EBs at 56°C for 30 min. High protein binding ELISA plates (Costar) were coated with 1×10^6^ HKEBs and blocked before serial dilutions of serum samples were added to the plates. *Chlamydia*-specific Abs were detected using HRP-based SBA Clonotyping System (Southern Biotech).

### Multiplex cytokine/chemokine array

At 10 days post *C. muridarum* intravaginal infection, five million cells from DLNs and total cells from the FRT were cultured *ex vivo* with 1×10^6^/mL HKEBs for 72 hours. Culture supernatants were collected and analyzed using the Mouse Cytokine/Chemokine 31-Plex Discovery Assay Array (Eve Technologies, Canada).

### Flow cytometry

Spleen, DLNs, and FRT were collected between days 10-14 post infection, and single cell suspensions prepared in RPMI containing 5% fetal bovine serum (FBS). FRTs were digested with collagenase IV (Sigma) at 37°C by GentleMACS (Miltenyi Biotech), and live cells purified using a Percoll gradient. For intracellular cytokine staining, cells were stimulated with PMA (50 ng/mL) and ionomycin (500 ng/mL) for 3 hrs at 37°C prior to surface and intracellular staining using the BD Cytofix/Cytoperm Kit (BD Biosciences). Intracellular transcription factor staining was performed using a Foxp3 staining kit (ThermoFisher). The following anti-mouse antibodies were obtained from BioLegend: FITC anti-CD4 (RM4-5), BV785 anti-CD4 (RM4-5), BV605 anti-CD8a (53-6.7), APC/Fire 750 anti-CD11b (M1/70), APC anti-CD44 (IM7), PE anti-CD44 (IM7), 0PerCP/Cy5.5 anti-CD45.1 (A20), Alexa Fluor 700 anti-CD45.2 (104), PE anti-CD45.2 (104), APC/Fire 750 anti-CD45R (B220; RA3-6B2), Alexa Fluor 700 anti-CD90.1 (OX-7), PE-Cy7 anti-CD90.1 (OX-7), Alexa Fluor 700 anti-CD90.2 (53-2.1), APC/Fire 750 anti-CD90.2 (53-2.1), APC/Fire 750 anti-F4/80 (BM8), FITC anti-IFN-γ (XMG1.2), Alexa Fluor 647 anti-IL-10 (JES5-16E3) and PE anti-GM-CSF (MP1-22E9). The following anti-mouse antibodies were obtained from BD Biosciences: BV510 anti-CD44 (IM7) and Alexa Fluor 488 anti-IL-17A (TC11-18H10). The following anti-mouse antibodies were obtained from ThermoFisher (eBioscience): PerCP-eFluor 710 anti-CD4 (RM4-5), PE-Cy7 anti-CD44 (IM7), PE anti-TCRVβ10b (B21.5), eFluor 450 anti-IFN-γ (XMG1.2), and eFluor 450 anti-FOXP3 (FJK-16S). Flow cytometry data were collected on an LSR Fortessa (BD Biosciences), an LSR Celesta (BD Biosciences) or Northern Lights cytometer (Cytek Biosciences). Data were then analyzed using FlowJo software (BD Biosciences). Polyfunctional analysis was performed using SPICE 6 (Roederer et al., 2011).

### T cell adoptive transfer

CD4 T cells were isolated from the spleens of donor mice and purified using the mouse CD4 T Cell Isolation Kit (Miltenyi Biotec). The purity and cell count of donor CD4 T cells were assessed using flow cytometry. For experiments using WT and *Bhlhe40^−/−^* mice as donors, 3×10^6^ CD4 T cells from each donor were mixed at a 1:1 ratio and transferred intravenously into TCRβ^−/−^ recipient mice. For TP1 adoptive transfer, 5×10^4^ WT TP1 cells (CD90.1+) and 5-7.5×10^4^ *Bhlhe40^−/−^* TP1 cells (CD90.2+) were isolated, mixed and transferred intravenously into CD45.1 recipient mice. All recipient mice were challenged intravaginally with *C. muridarum* one day after adoptive transfer.

### Monoclonal Ab treatment

*InVivo*MAb anti-mouse IL-10R blocking antibody (clone 1B1.3A) was obtained from BioXcell. Antibody treatment was performed by IP injection of 0.25 mg of anti-IL10R on days −1, 1, 4, 7, 10, 14, 17 and 21 following *C. muridarum* intravaginal infection.

### Single-cell RNA sequencing and TCR profiling

FRT cells from WT and *Bhlhe40^−/−^* mice were isolated 14 days post *C. muridarum* intravaginal infection and stained with surface markers. Live CD11b^−^F4/80^−^B220^−^CD90.2^+^CD4^+^CD44^hi^ CD4 T cells were sorted using FACSAria III (BD Biosciences), partitioned using a Chromium Controller (10x Genomics), and libraries prepared using Chromium Next GEM Single Cell 5’ Reagent Kits v2 (Dual Index) (10x Genomics). GEX and VDJ libraries were sequenced on Illumina NovaSeq 6000 at the sequencing depths of >25,000 reads/cell and >5,000 reads/cell, respectively. Data analysis was conducted using Seurat v.4.3 (Hao et al., 2021) and Monocle 3 (Trapnell et al., 2014) under R v4.3.1 environment. Contaminating non-CD4 T cells were cleaned up by filtering out cells without VDJ tags and/or with the following features: *Cd8a*, *Ncr1*. To avoid clustering based on T cell clonotypes, “*Trav*” and “*Trbv*” were removed from features when conducting unsupervised clustering using UMAP.

### Statistical analysis

Statistical analysis was performed by using an unpaired *t* test for normally distributed continuous-variable comparisons and a Mann-Whitney U test for nonparametric comparisons using Prism (GraphPad Software).

## Acknowledgements

We thank Drs. Brian Edelson (Washington U), Jason Stumhofer (UAMS) and Jie Sun (U of Virginia) for providing us with the *Bhlhe40^−/−^* and *Bhlhe40^fl/fl^-Cd4-Cre* mice. This study was supported by grants from the National Institutes of Health to LXL (AI139124 and GM103625) and LH (AI175738), and Careers in Immunology Fellowship from the American Association of Immunologists to MABM.

