## Supplemental Materials for "BHLHE40 drives protective polyfunctional CD4 T cell differentiation in the female reproductive tract against *Chlamydia*"

### Fig. S1

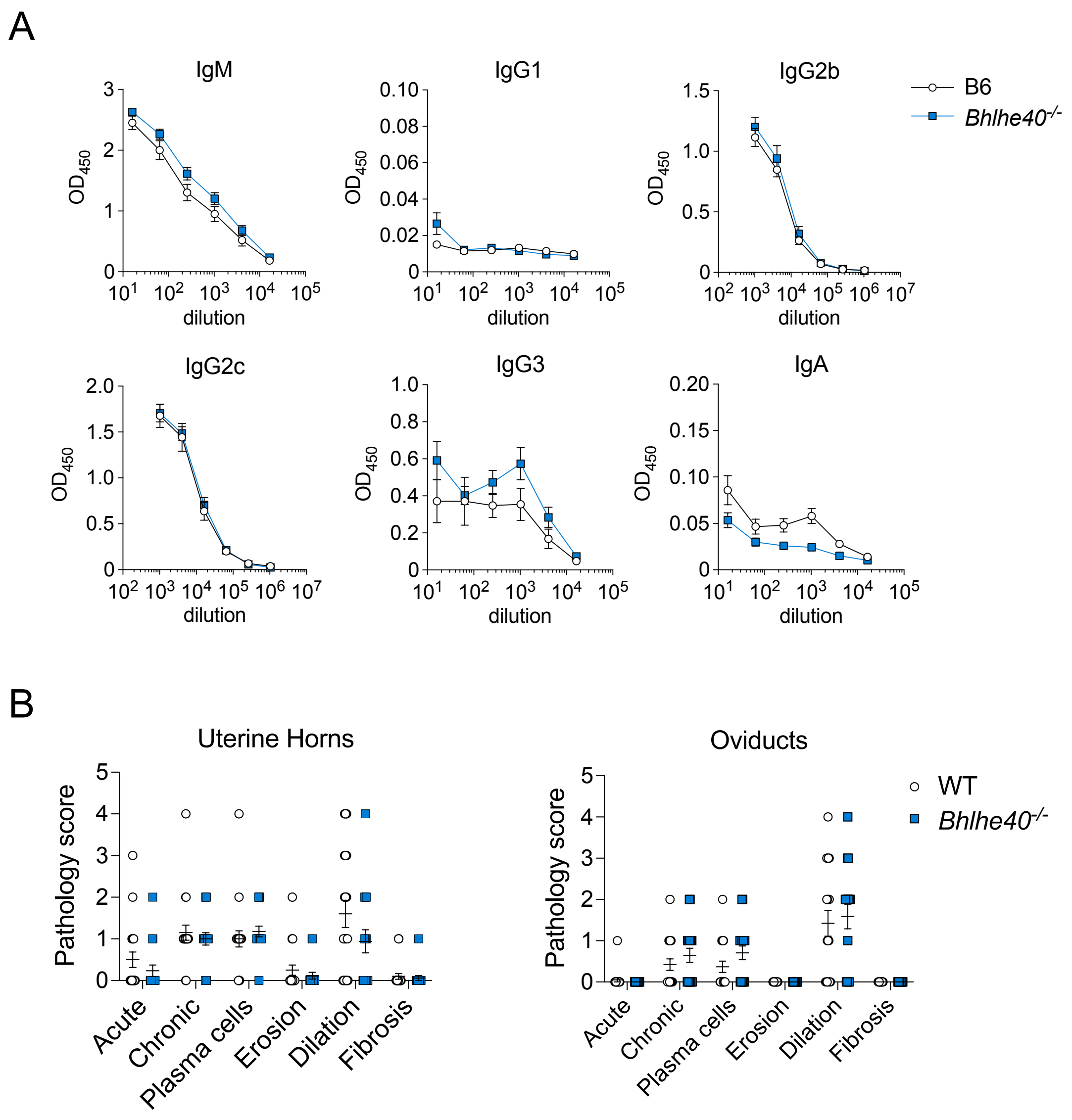


**Fig. S1. Antibody responses and FRT pathology are comparable between WT and *Bhlhe40*$^{-/-}$ mice.** WT and *Bhlhe40*$^{-/-}$ mice were infected with $1 \times 10^5$ *C. muridarum*.

**(A)** Anti-Cm serum antibodies were measured at day 21 post infection by EB Ab ELISA. Data are from two independent experiments with 9 mice per group. Error bars represent the mean ± SEM.

**(B)** FRTs were harvested between days 140-150 post infection. Pathology scores of uterine horns and oviducts were graphed for each category. Data are from two independent experiments with 8 to 10 mice per group. Each data point represents the left or right side of the corresponding section of the FRT. Error bars represent the mean ± SEM.

Fig. S2

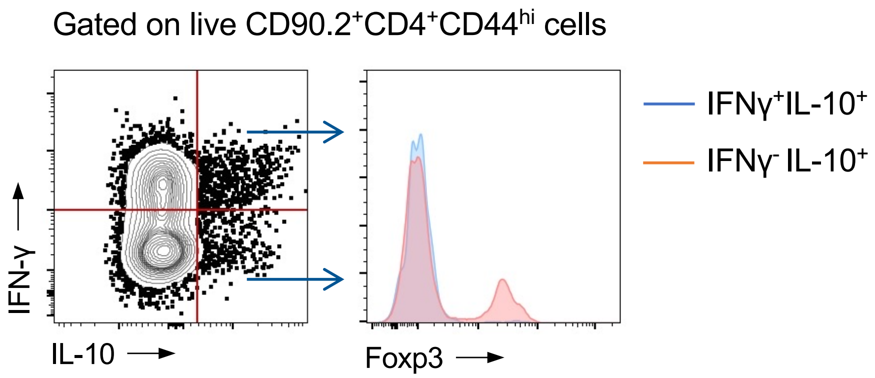


**Fig. S2. IFNγ+IL-10+ CD4 T cells were Foxp3- Tr1 cells.**
CD4 T cells from *Bhlhe40*-/- DLN samples in Fig. 2A were analyzed for Foxp3 expression by intranuclear staining. Data are representative of two independent experiments.

Table S1: Summary of cytokine multiplex data

| Cytokine | DLNs | | | | | FRT | | | | |
|---|---|---|---|---|---|---|---|---|---|---|
| | Concentration (pg/mL, ave ± SD) | | Ratio | *P* value (t test) | Significance | Concentration (pg/mL, ave ± SD) | | Ratio | *P* value (t test) | Significance |
| | WT (n=9) | *Bhlhe40* $^{-/-}$ (n=9) | WT/*Bhlhe40* $^{-/-}$ | | | WT (n=9) | *Bhlhe40* $^{-/-}$ (n=9) | WT/*Bhlhe40* $^{-/-}$ | | |
| Eotaxin | 2.1 ± 0.9 | 1.5 ± 1.1 | 1.4 | 0.23 | | 2.7 ± 1.1 | 11.4 ± 14.9 | 0.2 | 0.10 | |
| G-CSF | 577.8 ± 276.6 | 296.9 ± 239.5 | 1.9 | 0.035 | * | 6630.9 ± 2235.6 | 4190.1 ± 4168.5 | 1.6 | 0.14 | |
| GM-CSF | 147.5 ± 146.6 | 19.6 ± 12.0 | 7.5 | 0.019 | * | 494.6 ± 449.0 | 29.6 ± 18.3 | 16.7 | 0.0068 | ** |
| IFN-γ | 10476.3 ± 4471.3 | 2256.7 ± 1865.4 | 4.6 | 0.00011 | *** | 6970.4 ± 4250.0 | 111.6 ± 93.8 | 62.5 | 0.00018 | *** |
| IL-1α | 45.4 ± 29.6 | 23 ± 15.2 | 2.0 | 0.061 | | 224.7 ± 176.7 | 84.7 ± 69.9 | 2.7 | 0.042 | * |
| IL-1β | 27.9 ± 14.6 | 11.2 ± 3.3 | 2.5 | 0.0042 | ** | 275.8 ± 284.8 | 81.1 ± 68.1 | 3.4 | 0.063 | |
| IL-2 | 24.8 ± 14.8 | 19.4 ± 10.0 | 1.3 | 0.38 | | 13.3 ± 16.4 | 2.5 ± 1.3 | 5.2 | 0.067 | |
| IL-3 | 35.3 ± 36.7 | 11.1 ± 6.1 | 3.2 | 0.069 | | 34.2 ± 47.1 | 1.8 ± 0.7 | 18.9 | 0.055 | |
| IL-4 | 1 ± 1.3 | 0.4 ± 0.3 | 2.5 | 0.18 | | 2.2 ± 1.9 | 0.3 ± 0.2 | 5.9 | 0.023 | * |
| IL-5 | 506.6 ± 352.7 | 262.8 ± 89.2 | 1.9 | 0.062 | | 327.9 ± 163.4 | 117.3 ± 103.6 | 2.8 | 0.0049 | ** |
| IL-6 | 1697.1 ± 1025.7 | 380.2 ± 210.2 | 4.5 | 0.0017 | ** | 13855.9 ± 7916.9 | 8620.2 ± 11979.4 | 1.6 | 0.29 | |
| IL-7 | 1.8 ± 0.7 | 2.1 ± 0.9 | 0.8 | 0.40 | | 2.7 ± 1.3 | 3 ± 1.4 | 0.9 | 0.70 | |
| IL-9 | 16.7 ± 4.4 | 17.9 ± 3.5 | 0.9 | 0.53 | | 14.4 ± 6.7 | 12.9 ± 6.4 | 1.1 | 0.63 | |
| IL-10 | 180.5 ± 44.0 | 333.3 ± 267.4 | 0.5 | 0.11 | | 167.9 ± 101.4 | 129 ± 106.6 | 1.3 | 0.44 | |
| IL-12p40 | 3.7 ± 2.4 | 3.4 ± 1.5 | 1.1 | 0.78 | | 2.4 ± 0.8 | 1.9 ± 1.6 | 1.3 | 0.42 | |
| IL-12p70 | 8.2 ± 5.7 | 5.6 ± 3.1 | 1.4 | 0.26 | | 10.1 ± 5.9 | 8.2 ± 4.0 | 1.2 | 0.42 | |
| IL-13 | 0 | 0 | | | | 0 | 0 | | | |
| IL-15 | 8.9 ± 5.6 | 9.5 ± 5.3 | 0.9 | 0.80 | | 16.2 ± 7.8 | 15.6 ± 7.6 | 1.0 | 0.88 | |
| IL-17 | 689.4 ± 569.6 | 14.6 ± 7.7 | 47.0 | 0.0026 | ** | 901.9 ± 453.2 | 46.1 ± 46.9 | 19.5 | 0.000037 | *** |
| IP-10 | 104.3 ± 64.9 | 85.5 ± 37.4 | 1.2 | 0.46 | | 2906.4 ± 3378.4 | 984.6 ± 720.4 | 3.0 | 0.11 | |
| KC | 741.4 ± 917.6 | 124.3 ± 67.3 | 6.0 | 0.061 | | 9194.9 ± 2919.8 | 6598.7 ± 4383.3 | 1.4 | 0.16 | |
| LIF | 33.9 ± 16.9 | 27.8 ± 12.1 | 1.2 | 0.40 | | 93.3 ± 28.6 | 54.3 ± 43.0 | 1.7 | 0.038 | * |
| LIX | 205.1 ± 95.1 | 96.4 ± 62.9 | 2.1 | 0.011 | * | 573.6 ± 280.1 | 509.5 ± 480.1 | 1.1 | 0.73 | |
| MCP-1 | 231 ± 143.8 | 626.1 ± 278.2 | 0.4 | 0.0016 | ** | 7119.7 ± 4963.6 | 6823 ± 4542.4 | 1.0 | 0.91 | |
| M-CSF | 2.1 ± 0.8 | 1.2 ± 0.4 | 1.7 | 0.0086 | ** | 7.7 ± 10.1 | 3.6 ± 1.8 | 2.1 | 0.25 | |
| MIG | 145.2 ± 80.8 | 90.9 ± 53.8 | 1.6 | 0.11 | | 472.1 ± 261.9 | 142.8 ± 167.4 | 3.3 | 0.0058 | ** |
| MIP-1α | 244.1 ± 335.2 | 427.1 ± 306.0 | 0.6 | 0.24 | | 437.1 ± 275.1 | 373.8 ± 159.2 | 1.2 | 0.56 | |
| MIP-1β | 891.4 ± 760.7 | 1882.7 ± 905.8 | 0.5 | 0.023 | * | 1640.2 ± 850.2 | 1461.7 ± 760.2 | 1.1 | 0.64 | |
| MIP-2 | 7664.4 ± 5288.1 | 1365.1 ± 865.6 | 5.6 | 0.0028 | ** | 13499.4 ± 6547.8 | 7021.6 ± 4415.2 | 1.9 | 0.026 | * |
| RANTES | 31.8 ± 13.6 | 25 ± 6.1 | 1.3 | 0.19 | | 79.4 ± 18.6 | 49.9 ± 30.0 | 1.6 | 0.023 | * |
| TNFα | 45.1 ± 29.8 | 27.7 ± 9.0 | 1.6 | 0.11 | | 142.1 ± 85.6 | 40.3 ± 14.8 | 3.5 | 0.0029 | ** |
| VEGF | 290.7 ± 279.1 | 78.4 ± 34.2 | 3.7 | 0.038 | * | 495.9 ± 206.3 | 330.4 ± 187.5 | 1.5 | 0.094 | |

*, $p < 0.05$; **, $p < 0.01$; ***, $p < 0.001$
